# Modelling the efficacy of Neprilysin from various species in degrading different Amyloid-β peptides: Potential application in therapeutics of Alzheimer’s disease

**DOI:** 10.1101/2021.03.25.436981

**Authors:** Arun HS Kumar

## Abstract

Recombinant neprilysin due to its degradation potential against Amyloid-β (Aβ) peptides has been looked at as a potential therapeutic candidate for treating Alzheimer’s disease (AD). However the enzymatic activity of neprilysin against different Aβ peptides can variable which significantly limits the therapeutic optimization. Using the molecular interaction analysis and modelling it against the known enzyme-substrate kinetics, this study developed a novel approach to predicting biosimilar enzyme-substrate kinetics. The known enzyme-substrate kinetics of human recombinant neprilysin with Aβ_1-40_ peptide was used as the prototype to assess the affinity and efficacy of various inter and intra-species neprilysin- Aβ peptide enzyme kinetics based on the relative molecular interaction analysis. Significant inter and intra-species variations in neprilysin- Aβ peptide enzyme kinetics was observed which further validated the need for optimizing enzyme kinetics tailored to specific substrate degradation. The novel enzyme kinetics modelling approach described in this study can be helpful in the developing of recombinant enzymes/peptides for personalised therapeutic applications.

## 1. Introduction

Neprilysin is a peptidase which is reported to cleave Amyloid-β (Aβ) peptides associated with Alzheimer’s disease (AD).^1–3^ Aβ peptides are major constituents of amyloid plaques which correlate with progression of AD in humans.^1–3^ Despite the well-known role of Aβ peptides in AD, the rodent models of AD, which have been the predominant tools for discovery of AD therapeutics don’t show considerable presence of amyloid plaques or the related neurofibrillary tangles.^4, 5^ This discrepancy has been attributed to species specific differences in the amino acid sequences of Aβ peptides which consequently influence enzymatic activity of neprilysin against Aβ peptides,^1, 4^ Further recent studies have suggested that neprilysin can degrade murine Aβ with a much higher efficiency than the human Aβ,^1^ which may partly explain the reasons for failure to observe amyloid plaques in the rodent models of AD. The Aβ are produced as a consequence to degradation of amyloid precursor protein (potentially by neprilysin).^3, 6–8^ Several soluble forms of Aβ are produced which may have a variable enzyme-substrate kinetics with the neprilysin.^3, 6–8^ Likewise six potential isoforms of neprilysin are also reported of which only two isoforms have been cloned and are reported to have high degree of homology.

Considering the efficiency of neprilysin to cleave Aβ peptides, there may be merit in administering stabilised forms of neprilysin as a therapeutics for AD.^9, 10^ However development of neprilysin to deliver it to its site of action in the brain and achieve required therapeutic efficacy comes with several challenges. One such challenge is the effectiveness with which various forms of Aβ can be cleaved to facilitate their optimal removal from the brain. The cleaving of Aβ peptides to achieve an optimal physiological balance is essential as neprilysin knockout mice are reported to have better cognitive potential with aging,^1^ suggesting neither neprilysin inhibitors nor its activator in exclusivity can be therapeutically beneficial in AD. Under these circumstances controlled delivery of recombinant neprilysin for optimal regulation of Aβ peptides in the brain will be therapeutically beneficial in AD. Developing recombinant neprilysin for therapeutic use will require a detailed understanding of the enzyme kinetics between neprilysin and Aβ peptides. Hence this study modelled the enzyme kinetics between different forms neprilysin and Aβ peptides, which can be helpful in optimization of recombinant neprilysin for therapeutic development.

## 2. Materials and Methods

### 2.1. Protein structure

The protein data bank (PDB; https://www.rcsb.org/) was searched with Neprilysin and Amyloid β peptides as key words to identify the reported 3D structures. There were five Neprilysin and multiple Amyloid β peptide 3D structures identified in the PDB database. Of the five Neprilysin structures, two each were of human and bacterial origin, while one was from rabbits. The following neprilysin and Amyloid β peptides (PDB ID given) were used: 5JMY (Human Neprilysin from Spodoptera frugiperda expression system, 6GID (Human Neprilysin from Komagataella pastoris expression system), 4XBH (Soluble rabbit neprilysin), 6GHX (Bacterial Thermolysin; Geobacillus stearothermophilus), 1THL (Bacterial Thermolysin; Bacillus thermoproteolyticus), 4XFO (Amyloid-forming segment TAVVTN from human Transthyretin), 5TPT (Amyloid Precursor-Like Protein 2 (APLP2) E2 Domain;Human), 3SGP (Amyloid segment of alphaB-crystallin residues 90-100;Human), 2ROZ (Cytoplasmic tail of amyloid precursor protein;Mouse), 2YT1 (Cytoplasmic tail of amyloid precursor protein; Mouse), 4YN0 (Crystal structure of amyloid precursor protein E2 domain; Mouse), 2KJ7 (Rat Islet Amyloid Polypeptide), 1NMJ (Rat Amyloid beta-1-28), 4DBB (Amyloid precursor protein;Rat), 2BFI (Synthetic amyloid fibril), 2ONX (Amyloid cross-beta spines;Yeast) Additionally the reported sequence of human Amyloid β1-40 was used to construct the 3D structure using the Chimera software ^11–13^.

### 2.2. Molecular docking of the 3D structure

The 3D structure of Aβ peptides were individually docked against the five neprilysin structures (5JMY, 6GID, 4XBH, 6GHX and 1THL) to identify the number of interaction sites through formation of hydrogen bonds using the Chimera software.^11–13^ The number of interaction sites observed between the 5JMY and Amyloid β1-40 was taken as the baseline for all further estimations.

### 2.3. Enzyme kinetics modelling

The enzyme kinetics of 5JMY and Amyloid β1-40 interactions are extensively reported in the literature and forms the basis of several commercially available kits to assess neprilysin activity.^1, 6, 8, 14–16^ Hence the enzymatic cleaving of Amyloid β1-40 (at a substrate concentration of 0.625, 1.25, 2.5, 5, 10, 15 and 20 μM) was studied in presence of 5 nM concentration of recombinant neprilysin (5JMY). The resultant enzyme-substrate kinetic curve was used for modelling the kinetics of rest of the Amyloid β (4XFO, 5TPT, 3SGP, 2ROZ, 2YT1, 4YN0, 2KJ7, 1NMJ, 4DBB, 2BFI, 2ONX, Aβ1-40) and neprilysin (5JMY, 6GID, 4XBH and 6GHX) combinations. For the modelling of the enzyme-substrate kinetic curve, both a 3^rd^ and 4^th^ degree polynomial equation was used as these curves had high coefficient of determination i.e., r^2^=0.97 and 0.99 respectively, thus allowing better predictive power. In similar lines the enzyme kinetics was also modelled with fixed Amyloid β1-40 (15 μM) concentration and varying neprilysin (0.625, 1.25, 2.5, 5, 10, 15 and 20 ng) concentration. However in this case a second degree polynomial curve was used as this had a high coefficient of determination (r^2^=0.99). From the enzyme kinetic curves, a Lineweaver–Burk plot was made for each of the neprilysin and Amyloid β combinations to estimate the Vmax and Km values. Further to get an overview of the spread of Vmax and Km values for all the neprilysin and Amyloid β combinations, an innovative approach of plotting Km against the LogVmax was adopted in this study.

## 3. Results

The number of molecular interactions between different neprilysin and Amyloid β combinations was highly variable (Table 1). There were 48 hydrogen bonds observed between 5JMY and Amyloid β1-40 interactions. As this enzyme-substrate combination is widely studied in the literature, the number of hydrogen bond interactions in this combination was considered as baseline and formed the basis for modelling the enzyme-substrate kinetics of rest of the neprilysin and Amyloid β combinations. Accordingly the Michaelis–Menten kinetics of neprilysin (5JMY) and Amyloid β1-40 as a baseline was modelled to generate the enzyme saturation kinetics of rest of the combination based on the ratio differences (Hydrogen bond factor) in their respective hydrogen bonds shown in table 1. The Hydrogen bond factor was multiplied with the baseline curve generated either at constant neprilysin (Figure 1) or at constant Amyloid β (Figure 2) concentration. The considerable variations in the enzymatic activity of the different neprilysin and Amyloid β combination is evident from these graphs (Figure 1 and 2). All the four neprilysin evaluated in this study had greater affinity and efficacy towards the mouse Aβ peptides compared to human, rat, synthetic or yeast derived Aβ. The least enzymatic activity of neprilysin was observed against the yeast and synthetic Aβ.

**Figure 1.**
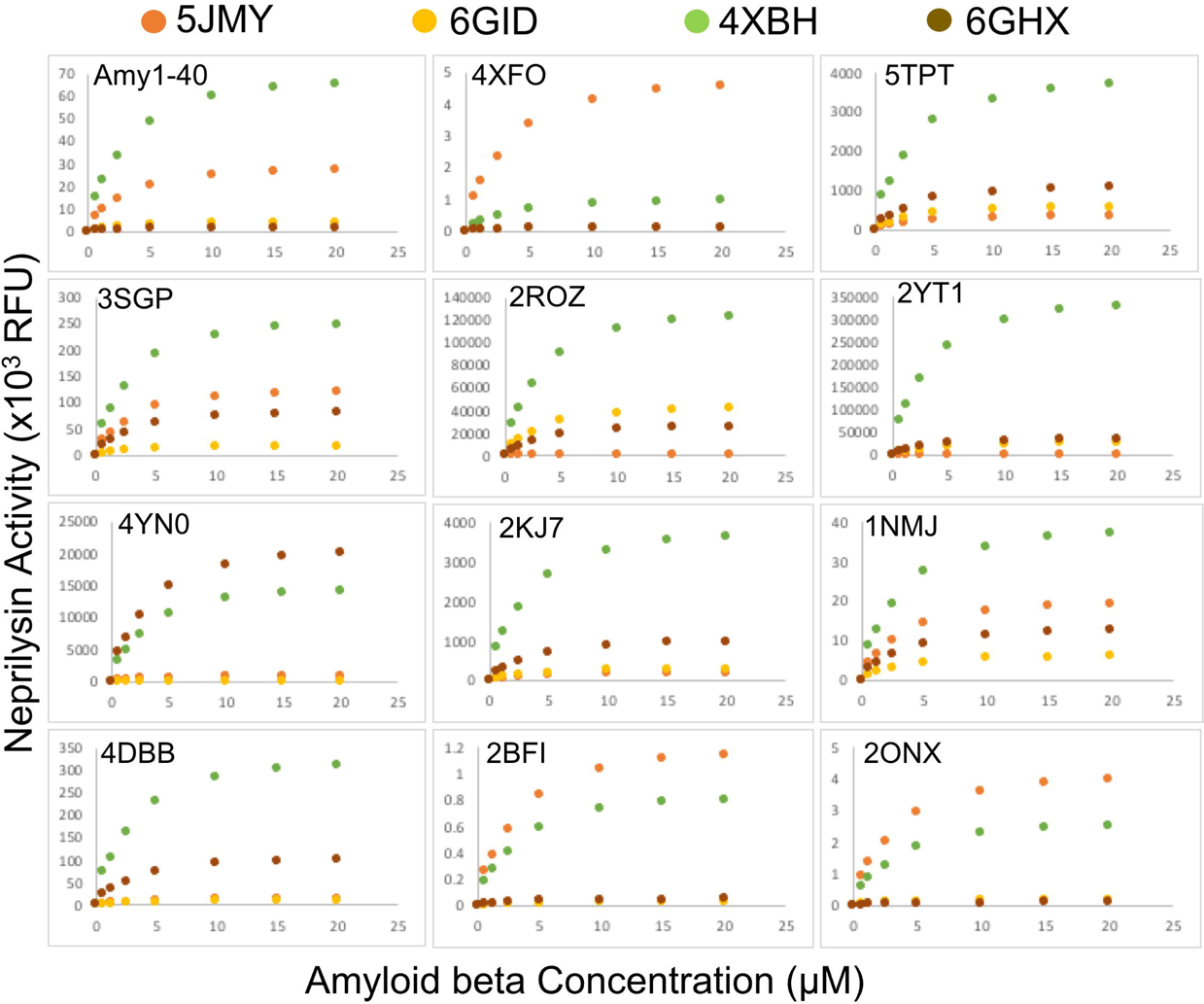
Enzyme-substrate kinetic curve of various neprilysin (5JMY, 6GID, 4XBH and 6GHX) and Amyloidβ peptides (4XFO, 5TPT, 3SGP, 2ROZ, 2YT1, 4YN0, 2KJ7, 1NMJ, 4DBB, 2BFI, 2ONX, Aβ_1-40_) under fixed neprilysin (5 nM) concentration.

**Figure 2.**
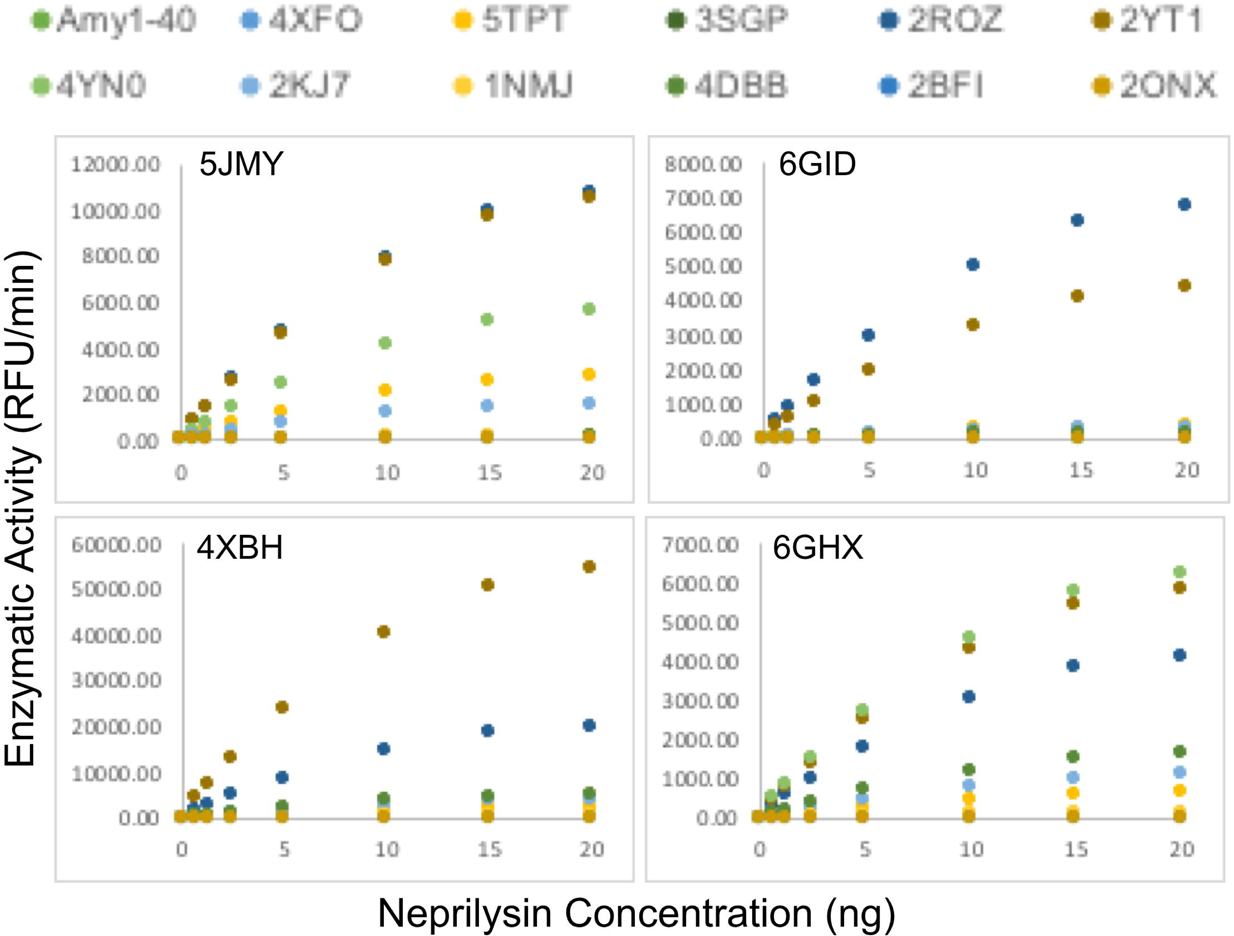
Enzyme-substrate kinetic curve of various neprilysin (5JMY, 6GID, 4XBH and 6GHX) and Amyloidβ peptides (4XFO, 5TPT, 3SGP, 2ROZ, 2YT1, 4YN0, 2KJ7, 1NMJ, 4DBB, 2BFI, 2ONX, Aβ_1-40_) under fixed substrate (15 μM) concentration.

Despite the considerable variations in the enzyme kinetics curves of various neprilysin and Aβ, the Km value of this reaction was consistently around 2.3±0.4 μM (Figure 3). The outlier values were mostly observed for Aβ peptides which had considerable isoforms (2Yt1, 2KJ7) or for the bacterial neprilysin. The affinity and efficacy of human neprilysin was specifically evaluated against the human Aβ verities (Table 2). The two recombinant human neprilysin assessed in this study were produced from different expression systems. Despite considerably homology between 5JMY and 6GID, the affinity (Km values) and efficacy (Specific Activity) of enzyme kinetics against different human Aβ peptides varied greatly (Table 2). 6GID-4XFO and 5JMY-4XFO enzyme kinetics showed the least Specific Activity and Km values respectively (Table 2). While the 6GID-5TPT and 6GID-4XFO enzyme kinetics showed the highest Specific Activity and Km values respectively (Table 2).

**Figure 3.**
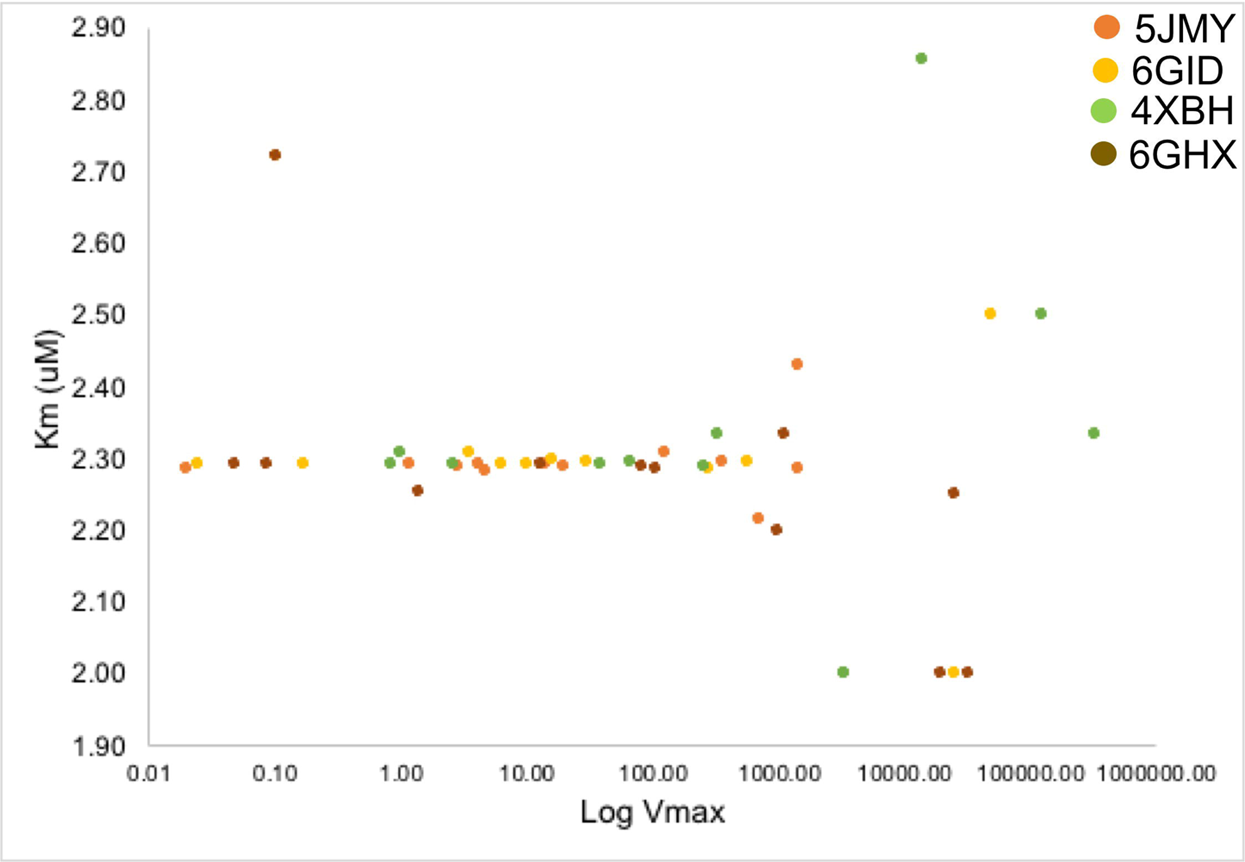
An XY plot of Km and Log Vmax values of various neprilysin (5JMY, 6GID, 4XBH and 6GHX) and Amyloidβ peptide (4XFO, 5TPT, 3SGP, 2ROZ, 2YT1, 4YN0, 2KJ7, 1NMJ, 4DBB, 2BFI, 2ONX, Aβ_1-40_) combination kinetics.

## 4. Discussion

This study reports a novel approach to evaluating enzyme-substrate kinetics based on the kinetics of a known prototype and estimating the kinetics of the novel enzyme-substrate types using the number of molecular interactions observed between the enzyme and substrate. Such a molecular modelling approach to predicting the enzyme-substrate kinetics can be valuable in the optimization of correct recombinant enzyme candidates for therapeutic development. This study also demonstrates the merit of the molecular modelling approach in predicting the enzyme-substrate kinetics using the neprilysin-Amyloidβ (Aβ) peptides as a prototype. The neprilysin- Amyloidβ (Aβ) was selected as the enzyme-substrate prototype as recent studies have indicate the potential role of developing recombinant neprilysin as a therapeutic candidate for treating Alzheimer’s disease (AD).^2, 6, 8, 9, 16^

Therapeutic development of neprilysin for use in treating AD will require establishing the optimal enzyme kinetics against AD specific Aβ peptides.^2, 8, 9^ Although the association Aβ peptides in the progression of AD is well established, developing effective treatment will require specific and selective cleaving of a variety of Aβ peptides.^14–17^ Depending on the variation in the site of cleavage in the amyloid precursor protein a variety of Aβ peptides can be generated, which may impact the pathophysiology of AD differently.^17–19^ The variable nature of Aβ peptides will influence the extent to which they are further enzymatically degraded by neprilysin. As shown in this study considerable variation in the affinity and efficacy of different recombinant human neprilysin against Aβ peptides was observed. Consistent with this observation a previous study has reported the significant variability in the efficiency with which neprilysin can degrade murine versus human Aβ peptides.^1, 3, 9^ Besides such inter-species variability in the neprilysin efficacy, this study also demonstrated prevalence of intra-species variability in humans. This intra-species variability will necessitate selection of optimal combination of different recombinant neprilysin for reducing Aβ peptides in AD. The approach described in this study provides an evidence based rationalization for developing such recombinant neprilysin combinations. It is necessary that the recombinant neprilysin combinations are aligned to the nature of Aβ peptides in the patients with AD and this can be achieved through current imaging modalities with spectral analysis feature.^10, 17–19^ Once the nature of Aβ peptides is established the modelling approach described in this study can be used to identify the recombinant neprilysin which will be most effective to optimally reduce Aβ peptides to achieve the desired therapeutic efficacy. For instance in this study although both human recombinant neprilysin (5JMY and 6GID) had similar efficiency to cleave Aβ_1-40_, they had variable efficiency against human Aβ precursor protein and Aβ peptide fragments. Hence suggesting that in the AD patients with higher levels of Aβ precursor protein, 6GID will have better efficacy than 5JMY. In contrast AD patients with higher levels of Aβ peptide fragments will benefit from 5JMY compared to using 6GID. Further such an approach based on patient specific Aβ peptides will be pave way for personalised medicine.

## 5. Conclusions

In conclusion, this study describes a novel approach of correlating the molecular modelling of enzyme-substrate interactions with enzyme kinetics of the known prototype to predict the enzyme kinetics of similar other enzyme and substrate combinations. Such an enzyme kinetics modelling approach can be helpful in the developing of recombinant enzymes/peptides for personalised therapeutic applications.

## Declaration of Conflict of interest

none

## Acknowledgement

Research support from University College Dublin-Seed funding/Output Based Research Support Scheme (R19862, 2019), Royal Society-UK (IES\R2\181067, 2018) and Stemcology (STGY2708, 2020) is acknowledged.

## Notes

### Competing Interest Statement

The authors have declared no competing interest.

